# Transcriptomic changes in oligodendrocytes and precursor cells predicts clinical outcomes of Parkinson’s disease

**DOI:** 10.1101/2023.05.11.540329

**Authors:** Mohammad Dehestani, Velina Kozareva, Cornelis Blauwendraat, Ernest Fraenkel, Thomas Gasser, Vikas Bansal

**Affiliations:** German Center for Neurodegenerative Diseases (DZNE), Tuebingen, Germany; Department of Neurodegeneration, Hertie Institute for Clinical Brain Research, University of Tuebingen, Tuebingen, Germany; Department of Biological Engineering, Massachusetts Institute of Technology, Cambridge, MA, USA; Laboratory for Neurogenetics, National Institute of Health NIH, Bethesda, Maryland, USA

**Keywords:** Parkinson’s disease, oligodendrocytes, oligodendrocyte precursor cells, snRNA-seq, polygenic risk scores, PD symptoms

## Abstract

Several prior studies have proposed the involvement of various brain regions and cell types in Parkinson’s disease (PD) pathology. Here, we performed snRNA-seq on the prefrontal cortex and anterior cingulate regions from post-mortem control and PD brain tissue. We found a significant association of oligodendrocytes (ODCs) and oligodendrocyte precursor cells (OPCs) with PD-linked risk loci and report several dysregulated genes and pathways, including regulation of tau-protein kinase activity, regulation of inclusion body assembly and protein processing involved in protein targeting to mitochondria. In an independent PD cohort with clinical measures (681 cases and 549 controls), polygenic risk scores derived from the dysregulated genes significantly predicted Montreal Cognitive Assessment (MoCA)-, and Beck Depression Inventory-II (BDI-II)-scores but not motor impairment (UPDRS-III). We extended our analysis of clinical outcome prediction by incorporating three separate datasets that were previously published by different laboratories. In the first dataset from the anterior cingulate cortex, we identified a correlation between ODCs and BDI-II. In the second dataset obtained from the substantia nigra (SN), OPCs displayed notable predictive ability for UPDRS-III. In the third dataset from the SN region, a distinct subtype of OPCs, labeled OPC_ADM, exhibited predictive ability for UPDRS-III. Intriguingly, the OPC_ADM cluster also demonstrated a significant increase in PD samples. These results suggest that by expanding our focus to glial cells, we can uncover region-specific molecular pathways associated with PD symptoms.

---

Parkinson’s disease (PD) is the second most common neurodegenerative disorder. It is characterized by the pathologic aggregation of alpha-synuclein and its pathology is known to progress in a predictable spatiotemporal manner. The spread of the disease pathology commences in the olfactory bulb and the lower brainstem and moves through the substantia nigra pars compacta (SN) in the midbrain, eventually reaching the meso- and neocortical areas^1,2^. PD arises from a complex interplay of various factors, such as aging, genetic predisposition, and environmental factors. While the cause of PD is unknown in most cases, specific mutations in genes such as *LRRK2* and *GBA1* have been identified to significantly increase the risk of developing the disease, although likely by affecting different molecular pathways. This may be one reason why the impact of *LRRK2* and *GBA1* mutations on clinical presentations may differ^3^. In recent decades, transcriptome profiling has emerged as a preeminent methodology for exploring human pathologies at the molecular and cellular level. In PD, alpha-synuclein pathology has been shown to be associated with the transcriptional programs of various brain cell types, including neurons and glial cells^4^. Furthermore, although the degeneration of dopaminergic neurons (DA) in PD primarily occurs in the SN region, Lewy bodies can form in other brain regions, such as the limbic system and the prefrontal cortex^5^. Moreover, there has been no investigation into the relative impacts of *LRRK2* and *GBA1* risk genotypes on transcriptional programs across several brain regions of PD patients.

In this study, we performed single-nucleus RNA-sequencing (snRNA-seq) on the prefrontal cortex (PFC) and anterior cingulate (ACC) brain regions from the same individuals (2 *LRRK2* PD, 2 *GBA1* PD and 2 Healthy Controls) to identify neuronal and non-neuronal cell-type differences (Figure 1A and Table S1). After data cleaning and quality control (see Methods), 88,876 high-quality single nuclei were retained. The clustering of these high-quality nuclei identified 13 clusters covering major cell types in the brain, i.e. excitatory neurons (ExN), inhibitory neurons (InN), oligodendrocytes (ODCs), oligodendrocyte precursor cells (OPCs), microglia (MG), astrocytes (Astro) and vascular cells (Vas) (Figure 1B). While ∼50% of the total nuclei were annotated as ODCs, only 4.3% and 0.5% of nuclei accounted for OPCs and Vas cells, respectively (Figure 1C). On average, we obtained 7406 high-quality nuclei per sample ranging from 3.6% to 16.5% of the total nuclei (Figure 1C and Table S2). While the percentage of each cell type varied across the samples, no significant differences (see Methods) in cell type proportions were observed between brain regions or mutation groups (Figure 1D-E and Table S2). The clusters were annotated based on the expression of well-known cell-type markers (Figure 1F and Table S3). The top 5 markers of each cell type cluster are shown in Figure 1G.

**Figure 1:**
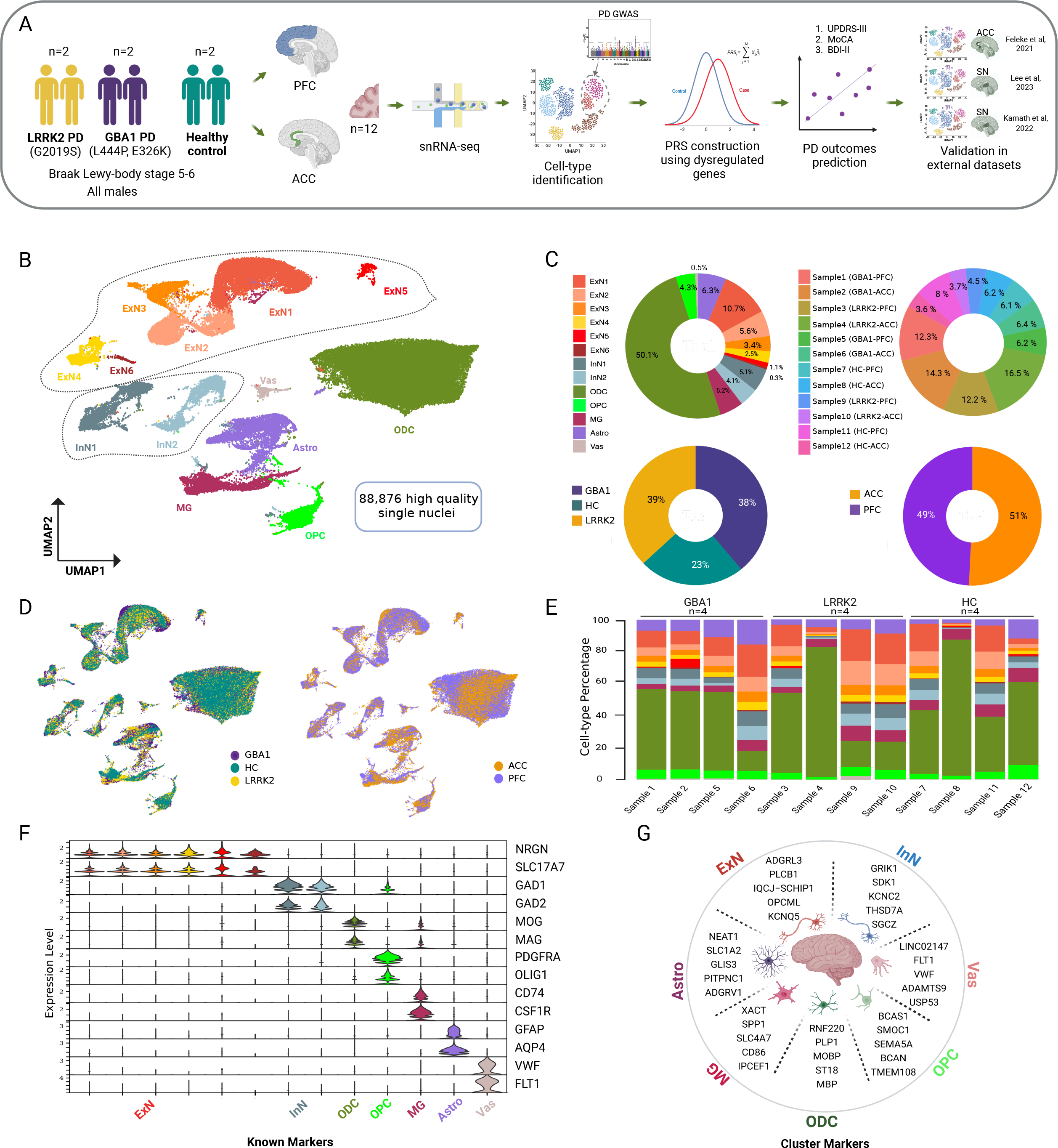
Overview of snRNA-seq profiling in the human post-mortem brain tissues. **(A)** Schematic overview of the experimental plan. **(B)** Uniform manifold approximation and projection (UMAP) visualization of the snRNA-seq clusters from 88,876 high quality nuclei. **(C)** Percentage of nuclei for each cell type across samples, mutation group and brain regions. **(D)** UMAP embeddings of nuclei colored by mutation group and brain regions. **(E)** Barplot displaying the distribution of cell-type percentage sample-wise. **(F)** Violin plot illustrating the expression distribution of known gene markers. **(G)** Genes most up-regulated in identified cell types: excitatory neurons (ExN), inhibitory neurons (InN), oligodendrocytes (ODCs), oligodendrocyte precursor cells (OPCs), microglia (MG), astrocytes (Astro) and vascular cells (Vas).

Integration of snRNA-seq data, which includes all expressed genes, with PD genome-wide association studies (GWAS)^6^ showed the strongest association of ODCs and OPCs with PD-linked risk loci (Figure 2A and Table S4). Next, we performed differential expression gene analysis to identify dysregulated genes between PD and controls in ODCs and OPCs. While 1040 and 543 genes were differentially expressed between *GBA1* and control samples in ODCs and OPCs, respectively, only 278 genes in ODCs and 108 genes in OPCs were differentially expressed between *LRRK2* and control samples (Figure 2B and Table S5). Using all the DEGs, MAGMA indicated that OPCs had the highest association with PD-linked risk loci (Figure 2C and Table S6). The most prominent MAGMA association was observed among DEGs in OPCs when comparing LRRK2 vs HC, aligning with earlier findings indicating highest *LRRK2* expression in OPCs^7^. Enriched biological processes in up-regulated DEGs (Figure 2D and Table S7) exhibited negative regulation of inclusion body assembly (in *GBA1*/Control ODCs), regulation of microtubule nucleation (in *LRRK2*/Control ODCs), positive regulation of tau-protein kinase activity (in *GBA1*/Control OPCs) and regulation of protein polymerization (in *LRRK2*/Control OPCs). On the other hand, enriched biological processes in down-regulated DEGs exhibited protein processing involved in protein targeting to mitochondria (in *GBA1*/Control ODCs), regulation of potassium ion transport (in LRRK2/Control ODCs), modulation of chemical synaptic transmission (in *GBA1*/Control OPCs) and cholesterol biosynthetic process (in *LRRK2*/Control OPCs).

**Figure 2:**
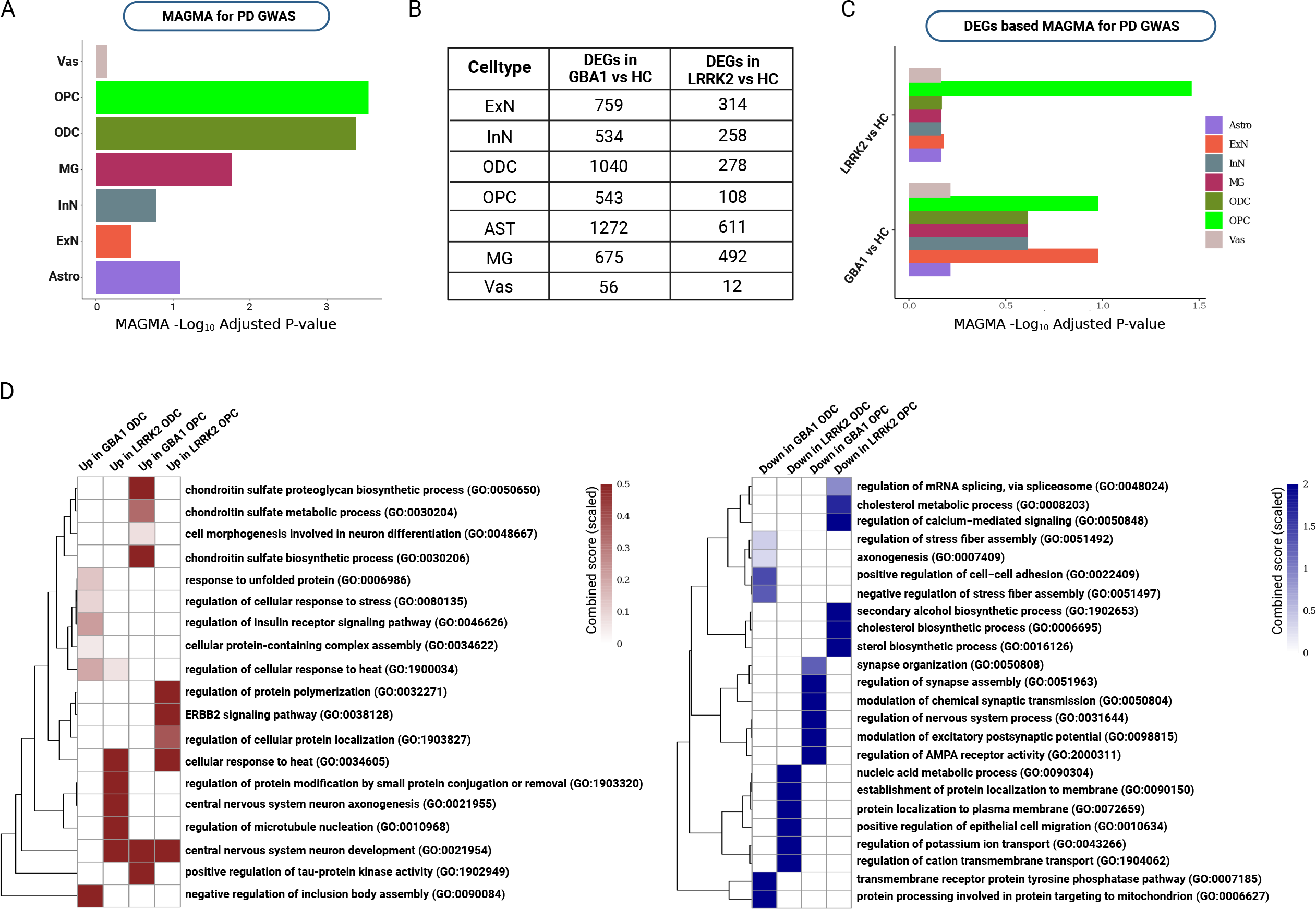
Association of PD susceptibility with ODCs and OPCs. **(A)** Multi-marker analysis of genomic annotation (MAGMA) gene set enrichment based on all the 88,876 high quality nuclei showed significant associations with oligodendrocytes (ODCs) and oligodendrocyte precursor cells (OPCs). **(B)** Number of differentially expressed genes (DEGs) in each comparison and cell-type. **(C)** MAGMA gene set enrichment based on DEGs in LRRK2 vs HC (upper) and in GBA1 vs HC (lower). **(D)** Gene ontology enrichment analysis of up-regulated (left) or down-regulated (right) genes. Top five biological process terms for each gene list are indicated. Enrichr combined score is calculated by the logarithmic transformation of the p-value obtained from Fisher’s exact test, multiplied by the z-score representing the deviation from the expected rank.

Next, we performed polygenic prediction in an independent PD cohort from Tuebingen (681 cases and 549 controls), which has a uniquely rich and detailed set of PD clinical measures. Polygenic Risk Scores (PRS) were calculated using the GWAS summary statistics for PD excluding the data from our Tuebingen cohort (see Methods). We computed four different PRS derived from four gene lists obtained through our analysis of differential expression in ODCs and OPCs (Table S8). All the scores significantly predicted case-control status (Table S8). Subsequently, we conducted prediction of PD measures among patients. We focused on Unified Parkinson Disease Rating Scale-III (UPDRS-III), Montreal Cognitive Assessment (MoCA) and Beck Depression Inventory-II (BDI-II) among PD patients (*N* ranges from 379 to 514). Significant associations were observed between GBA1_OPC_DEG score with MoCA, LRRK2_ODC_DEG and LRRK2_OPC_DEG scores with BDI-II. However, it is noteworthy that none of the scores were able to predict motor examination measures i.e. UPDRS-III (Table 1).

**Table 1:**
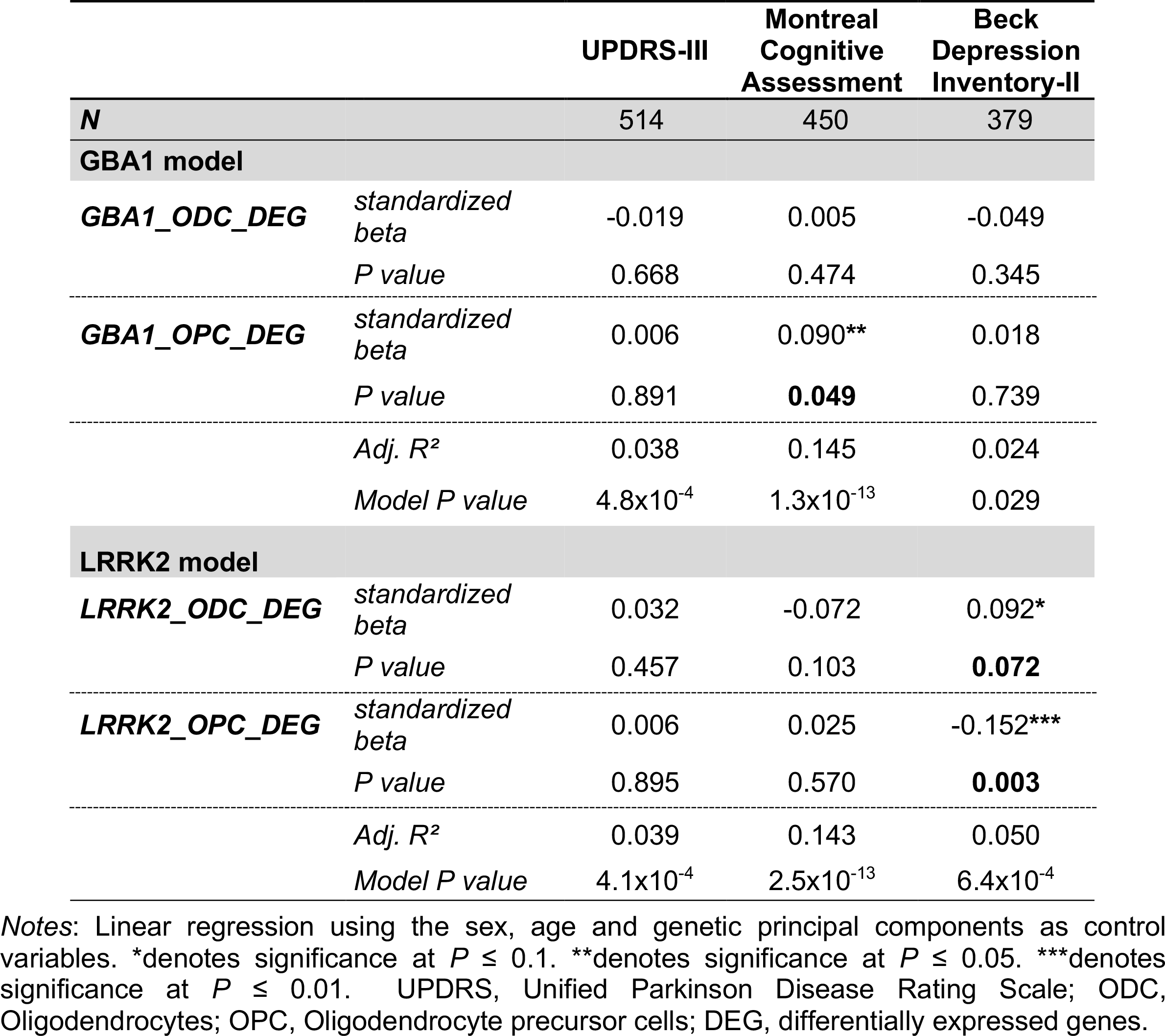
Polygenic prediction of Parkinson’s measures in the Tuebingen patient sample.

In order to confirm and broaden our findings beyond cortical regions, we utilized three distinct single-cell datasets for predicting PD measures using PRS derived from DEGs in ODCs and OPCs^8-10^ (Table S9). Using a dataset from the first study focusing on the ACC region^8^, we uncovered an association between ODC DEGs and BDI-II (Figure 3A and Table S10). In the second study centered on the SN region^9^, we discovered a substantial association between the OPC DEGs and UPDRS-III (Figure 3B and Table S10), indicating a potential specificity to different brain regions. To further uncover the association of PD measures and subtypes of OPCs and ODCs, we used snRNA-seq generated by Kamath and colleagues^10^, providing well-defined subtypes within the SN region. As a positive control, we first used DEGs from a highly vulnerable DA neuronal subpopulation, marked by SOX6_AGTR1. Indeed, as expected the PRS derived from DEGs in SOX6_AGTR1, were significantly associated with all three PD measures i.e. UPDRS-III, MoCA and BDI-II (Figure 3C and Table S10). Intriguingly, besides SOX6_AGTR1, we found a strong correlation of the OPC_ADM subtype with UPDRS-III (Figure 3C and Table S10). In addition, BDI-II was associated with the OPC_ADM, OPC_HOXD3 and ODC_ENPP6_EMILIN subtypes. It’s crucial to highlight that the OPC_ADM population showed a notable increase in PD samples (figure 3 of Kamath *et al*., 2022). Enriched biological processes in OPC_ADM DEGs exhibited distinct terms like regulation of myelination and glial cell differentiation whereas OPC_HOXD3 exhibited terms like regulation of receptor-mediated endocytosis, neuroblast proliferation and energy reserve metabolic process (Figure 3D and Table S11). ODC_ENPP6_EMILIN displayed terms related to unfolded protein and chaperone-mediated protein complex assembly (Figure 3D and Table S11). Similar processes have recently been found to be enriched in PD-associated oligodendrocytes^11^, suggesting that oligodendrocytes are affected by protein folding stress in PD.

**Figure 3:**
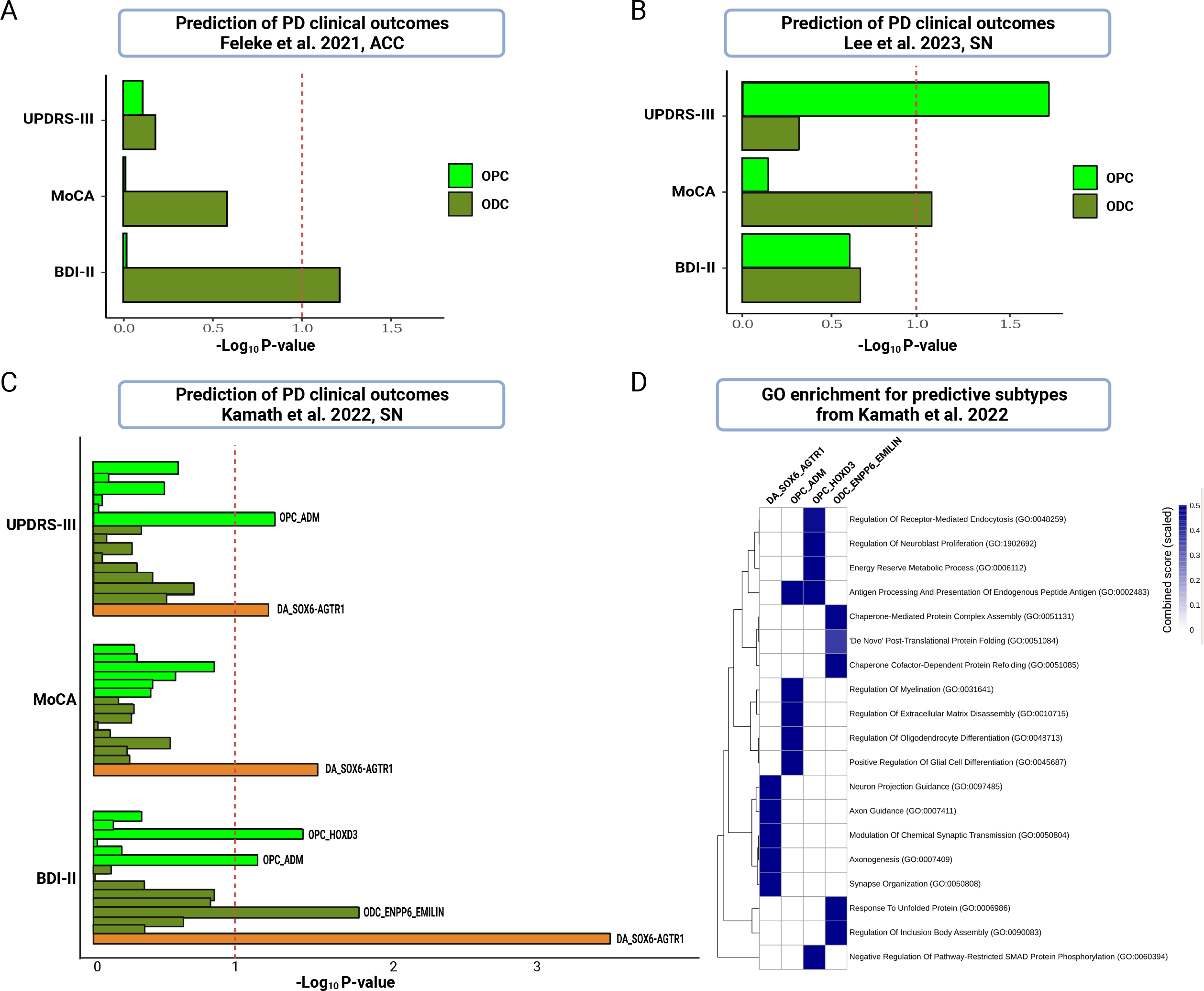
Polygenic prediction of PD measures using the ODCs and OPCs DEGs in publicly available datasets. **(A-C)** Prediction of clinical outcomes using Feleke *et al*., 2021 from anterior cingulate cortex region (A), Lee *et al*., 2023 from substantia nigra region (B) and Kamath *et al*., 2022 from substantia nigra region (C). **(D)** Gene ontology enrichment analysis of DEGs in predictive subpopulation of cell-types in Kamath *et al*., 2022 from substantia nigra region. Top five biological process terms for each gene list are indicated.

To summarize, in this short report, we found a significant association of PD GWAS risk loci in ODC and OPC expressed genes within PD GWAS risk loci and revealed several dysregulated genes and pathways, including regulation of tau-protein kinase activity, regulation of inclusion body assembly and protein processing involved in protein targeting to mitochondria. Accumulating evidence points out that oligodendrocytes and/or precursor cells also provide support to neurons via mechanisms beyond the insulating function of myelin^12-14^. Therefore, it is tempting to speculate that the abnormally regulated pathways, which extend beyond myelination, such as those involved in metabolic support to neurons, may contribute to the pathology of PD. Here, we would also address certain limitations of our study. We recognize that the sample sizes of the newly generated data in this study are comparatively small. Therefore, we utilized three distinct single-cell datasets from previously published studies to validate our findings. However, it is crucial to highlight that not all datasets exhibit uniform distribution in terms of age, gender, and mutations. Additionally, we are currently expanding our Tuebingen cohort to enhance the predictive power of clinical outcomes. In the future, it will be essential to integrate and replicate the results in a larger cohort characterized by a balanced metadata. It is noteworthy that in line with our results, previous studies indicate a significant enrichment of PD heritability in glial cell types like oligodendrocytes and astrocytes^15-17^. Two decades ago, Wakabyashi et al., observed an abnormal accumulation of alpha-synuclein in the oligodendrocytes within the substantia nigra of PD patients^18^. By integrating GWAS results with single-cell transcriptomic data, Bryois et al., and Agarwal et al., observed oligodendrocytes and oligodendrocyte precursor cells to be significantly associated with PD^16,17^. Moreover, while the loss of DA neurons in the SN region of the midbrain is a well-known pathological hallmark of PD closely associated with motor symptoms, it is important to note that PD patients also encounter various non-motor symptoms, including cognitive and psychopathological manifestations^19^. Szabolcs and colleagues found higher rates of psychiatric morbidity (especially mood disorders, cognitive impairment, anxiety disorders, schizophrenia) in the premotor phase of PD and these were more common in PD patients before PD diagnosis^20^. In line with this, the polygenic predictions in this study showed notable associations with non-motor symptoms, suggesting a crucial involvement of glial cells in neuropsychiatric symptoms that may extend beyond the SN region of the midbrain. On the other hand, significant correlation between OPC subpopulation in the SN region and PD motor symptoms implies region-specific alterations of molecular pathways in glial cells. Altogether, we anticipate that our study will serve as a valuable resource and prompt further research into the involvement of oligodendrocytes and oligodendrocyte precursor cells in the pathology of PD.

## Methods

### Samples used in this study

The research was conducted using fresh-frozen postmortem brain tissues obtained from four PD patients, two of which had the *LRRK2* p.G2019S mutation and the other two had the *GBA1* mutation (one with p.L444P and one with p.E326K), along with two healthy controls. All donors were males, aged between 65 and 80, and PD patients had Lewy body Braak stages of 5-6. Two brain regions, namely the prefrontal cortex and anterior cingulate cortex, were investigated for each donor. All tissues were procured from the Netherlands brain bank except one *LRRK2* brain from UCL Queen Square Brain Bank for Neurological Disorders, following the policies and regulations of the institutional ethics board at the University Hospital Tuebingen in Germany. For transcriptome analysis, single nuclei were extracted from all samples, and snRNA-seq was performed. The clinical characteristics of the donors have been elaborated in Table S1.

### Generation of single nuclei from postmortem human brains

The process of isolating nuclei entails the utilization of a detergent lysis technique, where a detergent is employed to break down the cellular membranes, followed by the centrifugal separation of the nuclei. In brief, 300 mg of post-mortem brain tissue was dounce-homogenized in 2 ml of Nuclei EZ Prep Lysis Buffer (Sigma Aldrich, MA, USA) spiked with 0.2 U µl-1 RNase inhibitor (Sigma Aldrich, MA, USA), 3.3 µl DTT (Thermo Fisher Scientific, MA, USA) and 33 µl of 10% Triton X100 which were added before incubating on ice for 5 minutes in a final volume of 10 ml. Homogenized tissue was washed with 3-4 ml washing buffer which was fresh PBSB 1% and then was filtered through a 70-µm cell strainer (BD Bioscience, NY, USA). Then using a long tube, 10 ml of 1.8 M ice-cold sucrose cushion solution is added to each sample i.e the roughly 3 ml lysate. After carefully and completely discarding the supernatant and the sucrose cushion layer containing debris and myelin, 1 ml PBS buffer added to resuspend the nuclei and 4 ml nuclei suspension buffer (1% BSA-PBS solution). It was finally centrifuged on 500 g for 5 minutes. At the end, 2 µl DAPI (Sigma Aldrich, MA, USA) with the concentration of 1:100 was added to stain the nuclei. Final centrifugation step on 500 g for 5 minutes was preceded by incubating the DAPI added suspension for 15-20 minutes in the cold & dark room on the rotation wheel at 4 C. Sorting buffer which consists of 99 µl PBS and 1 µl RNase inhibitor (Sigma Aldrich, MA, USA) was added to resuspend the nuclei and make them ready for quality/quantity inspection and then to run on a 10x genomic chromium controller. Quality assessment was performed using fluorescence-activated cell sorting (FACS) to detect all DAPI-positive events, i.e. individual nuclei comprising more than 95 % of all events.

### Droplet-based snRNA-seq using 10x Genomics

Single-nuclei suspension concentration was determined by automatic cell counting (DeNovix CellDrop, DE, USA) using an AO/PI viability assay (DeNovix, DE, USA) and counting nuclei as dead cells. Single-nucleus gene expression libraries were generated using the 10x Chromium Next gel beads-in-emulsion (GEM) Single Cell 3’ Reagent Kit v3.1 (10x Genomics, CA, USA) according to manufacturer’s instructions. In brief, cells were loaded on the Chromium Next GEM Chip G, which was subsequently run on the Chromium Controller (10x Genomics, CA, USA) to partition cells into GEMs. Cell lysis and reverse transcription of poly-adenylated mRNA occurred within the GEMs and resulted in cDNA with GEM-specific barcodes and transcript-specific unique molecular identifiers (UMIs). After breaking the emulsion, cDNA was amplified by PCR, enzymatically fragmented, end-repaired, extended with 3’ A-overhangs, and ligated to adapters. P5 and P7 sequences, as well as sample indices (Chromium i7 Multiplex kit, 10x Genomics, CA, USA), were added during the final PCR amplification step. The fragment size of the final libraries was determined using the Bioanalyzer High-Sensitivity DNA Kit (Agilent, CA, USA). Library concentration was determined using the Qubit dsDNA HS Assay Kit (Thermo Fisher Scientific, MA, USA). snRNA libraries were pooled and paired-end-sequenced on the Illumina NovaSeq 6000 platform (Illumina, CA, USA).

### snRNA-seq quality control

Samples were demultiplexed using Illumina’s bcl2fastq conversion tool and the 10x Genomics pipeline Cell Ranger count v6.0.1 to perform alignment against the 10x Genomics pre-built Cell Ranger reference GRCh38-2020-A (introns included), filtering, barcode counting, and UMI counting. As a default, a cut-off value of 200 unique molecular identifiers expressed in at least 3 cells was used to select nuclei of sufficient complexity for further analysis. Each sample’s count was normalized by the SCTransform method in Seurat v4.1.0^21^ with mitochondrial reads regressed out. Two approaches were combined for quality control: (1) Doublets and multiplets were filtered out using DoubletFinder v2.0.3^22^ for each individual sample; (2) outliers with a high ratio of mitochondrial and ribosomal counts (each >10%) and cells with low a number of genes (N < 1000) were removed. The core statistical parameters of DoubletFinder used to build artificial doublets for true doublet classification were determined automatically using recommended settings. After applying these filtering steps on 105,781 input nuclei, the dataset contained 88,876 high-quality single nuclei that were eligible for further analysis. We used the speckle R package v0.99.7 to analyze differences in cell type proportions^23^. We used the propeller function with CellType, SamplID and Mutation/Region columns from the Seurat MetaData object as input for clusters, sample and group, respectively.

### Cell annotations and differential expression

After combining all samples into a single Seurat object, genes were projected into principal component space using the principal component analysis (RunPCA). Harmony R package^24^ was used for integration as well as for removing unwanted effects across subjects. The first 12 PC dimensions of data processed with Harmony were used as inputs into the FindNeighbours, FindClusters (at 0.1 resolution obtained out of a range of tested resolutions (0.1, 0.2, 0.5, 1.0)) and RunUMAP functions of Seurat. In brief, a shared-nearest-neighbor graph was constructed on the basis of the Euclidean distance metric in principal component space, and cells were clustered using the Louvain algorithm. The RunUMAP function with default settings was used to calculate 2D UMAP coordinates and search for distinct cell populations. Cluster markers and differential expression testing was performed on Seurat “RNA” assay containing seurat log-normalized counts using default Wilcoxon method implemented in Seurat v4.1.0. Differential gene expression test between cases and controls was performed for each cell type using the Wilcoxon ranked sum method implemented within the FindMarkers function. Gene ontology enrichment analysis for biological processes was performed using EnrichR^25^. In addition, hierarchical clustering of enriched GO terms was performed for a set of paired comparisons, including *LRRK2* vs. HC and *GBA1* vs. HC brains, in which differentially expressed genes showed significant enrichment (Adjusted P-Value < 0.05).

### Cell-type association with genetic risk of PD

Association analysis of cell type-specific expressed genes with genetic risk of PD was performed as described previously^26^, using Multi-marker Analysis of GenoMic Annotation (MAGMA) v2.0.2, in order to identify disease-relevant cell types in the data^27,28^. MAGMA, as a gene set enrichment analysis method, tests the joint association of all SNPs in a gene with the phenotype, while accounting for LD structure between SNPs. Competitive gene set analysis was performed on SNP p-values from the latest PD GWAS summary statistics including 23andMe data and the publicly available European subset of 1000 Genomes Phase 3 was used as a reference panel to estimate LD between SNPs. SNPs were mapped to genes using NCBI GRCh37 build (annotation release 105). Gene boundaries were defined as the transcribed region of each gene. An extended window of 10 kb upstream and 1.5 kb downstream of each gene was added to the gene boundaries.

### Polygenic Risk Scores (PRS)

Using summary statistics data from PD meta GWAS (https://pdgenetics.org/resources), we performed a comprehensive annotation using the Region Annotation function in Annovar^29^ to find all genes corresponding to the SNPs in the whole genome and then generated the GWAS gene list. We performed an overlap analysis between the GWAS annotated genes and gene lists obtained from our differential expression comparisons in ODCs and OPCs. Next, we retrieved SNPs in each of the gene lists, including MAF, beta, and p-value from the base data summary statistics. For the prediction, we used imputed genotypes of 681 cases and 549 controls from the Tuebingen cohort^30^, which were not included in the base data. To construct polygenic risk score (PRS) models, we utilized the R package PRSice2 v2.3.5^31^. We applied clumping procedure using r2 > 0.1 and 1000 kb as the clumping parameters in PRSice2 and a p-value of 0.05 was chosen as the threshold to exclude non-significant SNPs. In other words, lead SNPs with a p-value of 0.05 from the LD-clumped list were included in the calculation of PRS used in the regression models. The null model is a logistic regression model that measures the power of covariates including age, sex, and genetic principal components (PC1-4) in the prediction of PD status, whereas the full model adds the PRS to the null model, thereby isolating the additive influence of the PRS on risk prediction. The R^2^ was also adjusted for an estimated PD prevalence of 0.005 on the liability scale. To predict the clinical measures, PRS were included in a linear regression model using the “lm” function in R. Standardized beta was obtained using the “lm.beta” function.

### Publicly available datasets used in this study

We used three published snRNA-seq datasets from human postmortem specimens: (1) Feleke *et al*., anterior cingulate cortex^8^ (n = 7 per group PD vs Controls, proportion female: control = 1/7, PD = 5/7), (2) Lee *et al*., substantia nigra^9^ (n = 6 PD vs n = 13 Controls, proportion female: control = 4/13, PD = 3/6), and (3) Kamath *et al*., substantia nigra^10^ (n = 7 PD vs n = 8 Controls, proportion female: control = 6/8, PD = 2/7). With the exception of Kamath *et al*., DEGs were directly obtained from the respective studies. DEGs between cases and controls from Kamath *et al*., were calculated using the Wilcoxon ranked sum method as implemented in Seurat’s FindMarkers function (Seurat v4.3.0.1). DEGs were computed for each cell type subpopulation based on subpopulations defined in Kamath et al. Filtered gene expression matrices and subpopulation annotations for Kamath et al. data were downloaded from Single Cell Portal (https://singlecell.broadinstitute.org/single_cell/study/SCP1768/) and converted into Seurat objects for log-normalization and differential gene expression analysis.

## Ethics approval and consent to participate

Our study was approved by the Ethics Committee of the University Hospital and Medical Faculty Tuebingen.

## Availability of data and materials

A Seurat object of the processed snRNA-seq data has been uploaded to Zenodo (https://doi.org/10.5281/zenodo.7886802).

## Competing interests

The authors declare that they have no competing interests.

## Funding

This work was funded by the Charitable Hertie Foundation, DFG grant (BA 7811 to V.B.) and in part by the Intramural Research Programs of the National Institute on Aging (NIA). V.B. is also supported by a Career Development Fellowship at DZNE Tuebingen.

## Authors’ contributions

M.D., T.G. and V.B. conceived the project. T.G. and V.B. supervised the study. M.D. performed all the computational analysis and nuclei isolation. V.K. and E.F. analyzed the publicly available datasets. C.B. and M.D. performed the MAGMA analysis. M.D. and V.B. wrote the manuscript. All authors contributed to and reviewed the manuscript.

## Acknowledgments

We thank the NGS Competence Center Tubingen core facility for library preparations and sequencing. We also thank the Jonas Neher research group at DZNE Tubingen for providing the protocols and FACS facility. We would like to express our gratitude to the brain banks and the donors. We also thank Manu Sharma, Claudia Schulte and Anastasia Illarionova for genotype data, clinical data and MAGMA script, respectively. BioRender was used to prepare figures.

## Supplementary table legends

**Table S1**: Sample overview with meta-data.

**Table S2**: Number of nuclei for cell types across samples, mutation group and brain regions.

**Table S3**: Known and identified cluster markers.

**Table S4**: MAGMA snRNA-seq cell-type enrichment results for PD GWAS.

**Table S5**: Differentially expressed genes in snRNA-seq for ODCs and OPCs.

**Table S6**: MAGMA snRNA-seq cell-type enrichment results for PD GWAS using DEGs.

**Table S7**: Enriched gene ontology terms for DEGs in ODCs and OPCs.

**Table S8**: Polygenic risk scores in Tuebingen PD cohort with case-control prediction.

**Table S9**: Differentially expressed genes in ODCs and OPCs of publicly available datasets.

**Table S10**: Polygenic prediction of PD measures using the ODCs and OPCs DEGs in publicly available datasets.

**Table S11**: Enriched gene ontology terms for DEGs in predictive subpopulation of cell-types in Kamath *et al*., 2022 from substantia nigra region.

## Notes

### Competing Interest Statement

The authors have declared no competing interest.

### Summary of Updates

In order to confirm and broaden our findings beyond cortical regions, we integrated three distinct single-cell datasets from previously published studies.

